# Riluzole reverses blood-testis barrier loss to rescue chemotherapy-induced male infertility by binding to TRPC

**DOI:** 10.1101/2024.04.26.591381

**Authors:** Rufei Huang, Huan Xia, Wanqing Lin, Zhaoyang Wang, Lu Li, Jingxian Deng, Tao Ye, Ziyi Li, Yan Yang, Yadong Huang

## Abstract

Cancer treatments, including cytotoxic therapy, often result in male infertility, necessitating the development of safe and effective strategies to preserve male reproductive potential during chemotherapy. Notably, our study uncovers the potential of repurposing riluzole, an FDA-approved drug for amyotrophic lateral sclerosis (ALS), in enhancing spermatogenesis. Hence, this research aims to explore the feasibility of utilizing riluzole to alleviate male infertility induced by busulfan (BSF), a commonly used chemotherapy drug. We established a BSF-induced oligospermia model in 4-week-old male mice and found that riluzole could effectively countered the detrimental effects of BSF on sperm production in mice with oligospermia. By restoring blood-testis barrier (BTB) functionality, riluzole improves sperm quality and reduces testicular atrophy. Through transcriptomic and molecular docking analyses, we identify transient receptor potential canonical subfamily member 5 (TRPC5) as a potential target for riluzole-mediated regulation of blood-testis barrier function. These findings propose riluzole as a promising therapeutic option for chemotherapy-induced male infertility, thereby addressing the fertility challenges associated with cancer treatments. Moreover, repurposing riluzole could streamline the drug development process, providing a cost-effective approach with reduced risk compared to developing entirely new drugs.

## Introduction

Infertility is a significant global health issue that affects approximately 9% to 20% of couples, with over 50% of cases attributed to male-related factors (Ávila, Vinay, Arese, Saso, & Rodrigo, 2022; Martínez et al., 2023). Abnormalities in spermatogenesis, influenced by factors such as genetics, hormonal disorders, psychological stress, sexual issues, obesity, and medication, are often the cause (Choy & Amory, 2020; Choy & Eisenberg, 2018). Chemotherapies, particularly those involving alkylating agents like busulfan (BSF), are associated with reversible or irreversible fertility disorders (Qu, Itoh, & Sakabe, 2019).

Busulfan, a common chemotherapy drug, is one of the few anti-cancer drugs used in children, particularly those under the age of three (Combarel, Tran, Delahousse, Vassal, & Paci, 2023; Veal et al., 2012). Although the promising survival rate of childhood cancers who may receive chemotherapies, their fertility is impaired when they enter reproductive age (Meistrich, 2009). In a cohort of more than 10 years of follow-up observation among 214 survivors of childhood cancers with alkylating agent chemotherapy, azoospermia was observed in 25% of patients, and oligospermia was observed in 28% (Green et al., 2014). The development of assisted reproductive technology (ART) has provided hope for severely oligospermic men to become fathers (Harel, Fermé, & Poirot, 2011; Mazzilli et al., 2023). Nonetheless, these methods are relatively costly and ineffective against azoospermia resulting from failed spermatogenesis (Dohle, 2010). Such impacts seriously affect post-treatment quality of life in cancer survivors. Therefore, it is an urgent matter to explore an efficient and safe approach to preserve male future reproductive capacity.

Spermatogonial stem cells (SSCs) are crucial for initiating and maintaining spermatogenesis, the process of sperm production in the testes (Diao, Turek, John, Fang, & Reijo Pera, 2022). This process begins at puberty and continues throughout a man’s life. Preserving SSCs is essential to ensure the potential for spermatogenesis recovery after cytotoxic therapy (Gouk, Loh, Kumar, Watson, & Kuleshova, 2011; Okada & Fujisawa, 2019). In testis, Sertoli cells (SCs) are important somatic cells that nourish and develop SSCs by providing nutrients, cytokines, and paracrine factors (Hai et al., 2014; Tao et al., 2021). Additionally, SCs form tight junctions (TJs) as part of the blood-testis barrier (BTB), which is necessary for normal spermatogenesis (Mruk & Cheng, 2015). The BTB prevents the passage of macromolecules between the basal and adluminal compartments and offers numerous essential conditions for spermatogenesis, including testosterone concentration, ion regulation, immune-privileged microenvironment, and physical barriers (Kaur, Thompson, & Dufour, 2014; N. Li, Tang, & Cheng, 2016; Q. Wen et al., 2018). Proteins such as Zonula occludens-1 (ZO-1), claudins, occludins and connexin 43 (Cx43) contribute to the formation, stability, and function of tight junctions. They play essential roles in controlling paracellular transport, maintaining barriers of BTB, and regulating spermatogenesis (M. W. Li, Mruk, Lee, & Cheng, 2010; Stanton, 2016; Su et al., 2020). The deficiency of these proteins can lead to spermatogenesis failure. Furthermore, SCs are the main targets of toxicants in the testis (Gao, Mruk, & Cheng, 2015). Previous study revealed that BSF impairs the seminiferous tubule structure, including the SCs integrity, leading to germ cells death (Chen, Liang, & Wang, 2018). Therefore, targeting the functionality of SCs and BTB may prove pivotal for addressing spermatogenic dysfunction and establishing new therapeutic approaches.

In our previous study, we accidentally found that riluzole, the first drug approved by the FDA for the clinical treatment of amyotrophic lateral sclerosis (ALS), had the ability to induce reprogramming of mouse embryonic fibroblastic cells (MEFs) into Sertoli-like cells (CiSCs) (Y. Yang, Li, et al., 2022). Interestingly, our study also revealed that treatment with riluzole resulted in a significant upregulation of gene expression levels of various secreted factors in SCs. These factors, including *Gdnf*, *Bmp4*, *Scf*, *Cxcl12*, *Inhibin B*, and *Fgf2,* are known to be critical for the proliferation and differentiation of SSCs (Hofmann & McBeath, 2022; Sofikitis et al., 2008). This observation suggested that riluzole had the potential to regulate spermatogenesis and may be a promising treatment option for male fertility preservation in clinic.

Here, we observed that mice treated with riluzole for a period of just 7 days following BSF injection exhibited substantial alleviation of cytotoxic effects, protecting spermatogonia from the BSF. This protective effect was achieved by restoring the tight junction function of SCs. These findings hold promise for the development of effective treatments and therapeutic strategies aimed at mitigating the detrimental impact of chemotherapy on male fertility.

## Results

### Riluzole improved fertility in BSF-treated mice by enhancing sperm quality and repairing damaged reproductive organs

To evaluate the therapeutic potential of riluzole in treating BSF-induced reproductive injury, the oligospermic mice induced by BSF (30 mg/kg·bw) received riluzole therapy (3 mg/kg·bw for intraperitoneal injection) once daily for 1 week. Following treatment, the mice were fed normally diets and monitored until 8th week (Figure 1A). Mice treated with riluzole did not show any significant changes in body weight (Supplementary file 1). Mating experiments were conducted, and the result revealed that out of the 6 female mice in each group for the mating experiment, only 1 mouse was pregnant in the BSF group. However, 4 mice were pregnant in the riluzole-treated group, approaching levels comparable to those observed in the control group (5 mice) (Figure 1B). Semen analysis conducted using the CASA system revealed that riluzole treatment displayed notable enhancements in multiple parameters. These improvements comprised elevated sperm counts (Figure 1C-D), enhanced sperm viability (Figure 1E), improved motility (Figure 1F), as well as augmented motion parameters (VSL, VCL, VAP, ALH, MAD, BCF, STR, and LIN) (Supplementary file 1) when compared to mice exposed to BSF without receiving any treatment. Additionally, riluzole treatment effectively reduced the rates of sperm abnormalities (Figure 1G).

**Figure 1.**
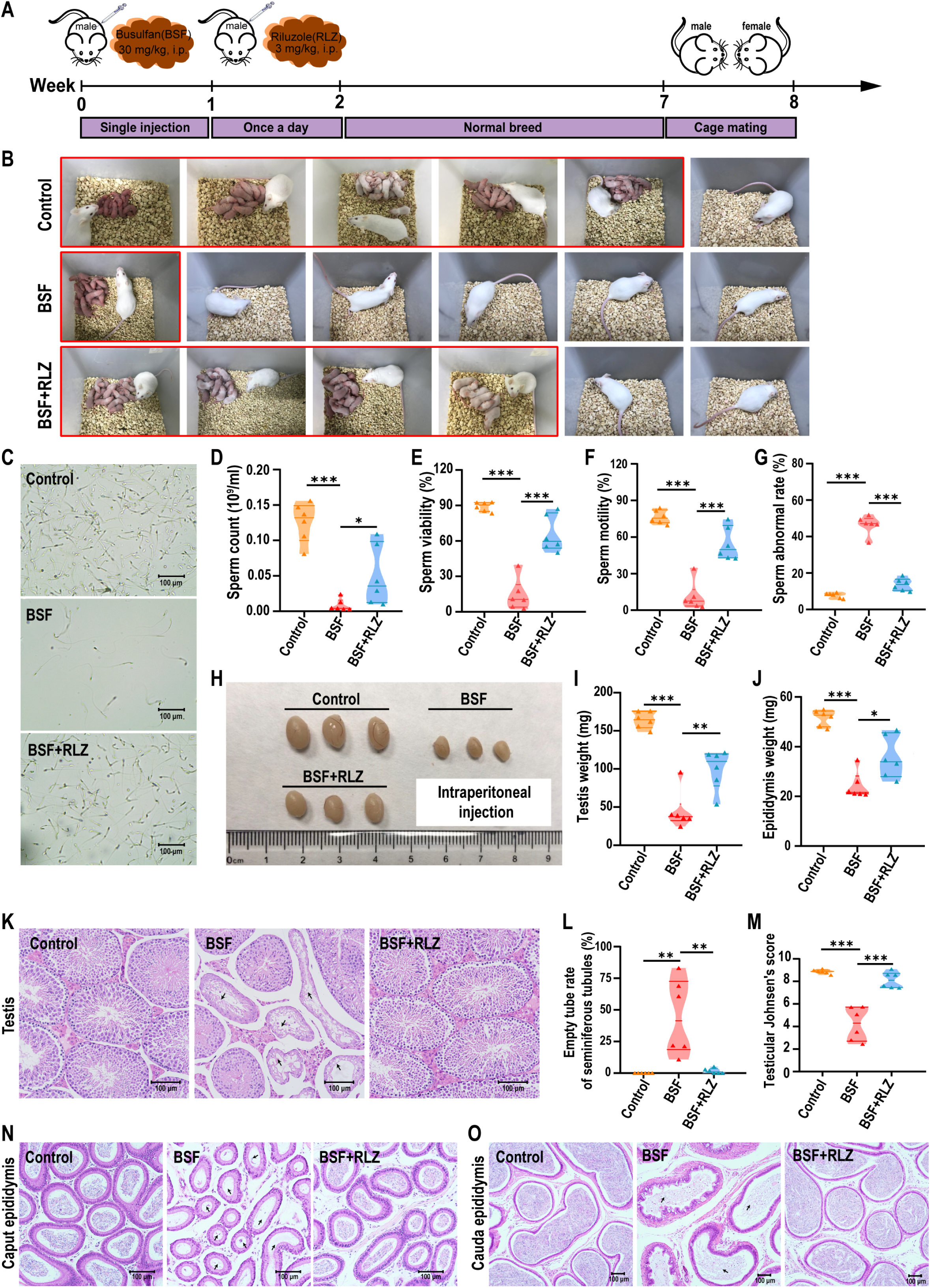
Effects of intraperitoneal administration of riluzole on sperm quality and recovery of damaged reproductive organs in oligospermic mice. (**A**) Illustration of experimental scheme. (**B**) Fertility of oligospermic mice induced by BSF after riluzole treatment (n=6). (**C**) Representative micrographs of sperms released from the cauda epididymis of BSF induced oligospermic mice after riluzole therapy. Scale bar, 100 μm. (**D-G**) Count, viability, motility, and abnormal rate of sperms in the cauda epididymis (n = 6). Sperm counts over 1000 were analyzed using CASA. (**H**) Morphology of the testes at week 8 from oligospermic mice after riluzole treatment. (**I-J**) Testis and epididymis weight of oligospermic mice after riluzole treatment (n =6). (**K**) Representative histopathology of the testes of mice with oligospermia after riluzole therapy. Black arrow presented low numbers of germ cells in the seminiferous tubule lumen. Scale bar, 100 μm. (**L**) Rate of empty tubes in seminiferous tubules after riluzole therapy (n=6). (**M**) Testicular Johnsen’s score of seminiferous epithelium (n=6). The seminiferous epithelium was evaluated according to the description of Johnsen’s score standard. (**N-O**) Representative histopathology of the caput epididymis and cauda epididymis of mice with oligospermia after riluzole therapy. Black arrow presented lower density of sperms in the epididymis. Scale bar, 100 μm. Values here are expressed as mean ± SEM. One-way analysis of variance (ANOVA) was used to analyze statistical differences; \*\*\**P* < 0.001, \*\**P* < 0.01, \**P* <0.05 compared to oligospermic mice treated with the vehicle (BSF).

Then mice were sacrificed to evaluated the effects of riluzole therapy on the male reproductive system. Results suggested that riluzole treatment successfully reversed testicular atrophy in oligospermic mice induced by BSF (Figure 1H). The weight of the testes and epididymis of mice treatment with riluzole significantly increased in comparison to oligospermic mice induced by BSF (testes: 99.99 ± 10.74 mg vs. 45.08 ± 10.40 mg; epididymis: 35.73 ± 3.48 mg vs. 24.30 ± 2.22 mg) (Figure 1I-J). Riluzole administration significantly reduced vacuolization induced by BSF and increased the number of germ cells in the seminiferous tubules (Figure 1K-L). Moreover, following riluzole treatment, the Johnsen score of the seminiferous epithelium (8.13 ± 0.29 vs. 4.21 ± 0.61) and the number of mature sperm in the epididymis was significantly increased (Figure 1M-O). These results suggested that riluzole had the potential to significantly improve fertility by enhancing sperm quality and rescuing damaged reproductive organs in mice with oligospermia.

### Intragastric gavage administration of riluzole rescued fertility in oligospermic mice induced by BSF

To address concerns about the potential damage to the abdominal viscera and improve patient compliance, we decided to switched to a safer method of administering riluzole. Instead of using intraperitoneal injection, we chose to administer riluzole through intragastric gavage (Figure 2A). Mice induced with oligospermia by the BSF were given riluzole through intragastric gavage once daily for 1 week. Following an 8-week period, mating experiments confirmed that, similar to the previous intraperitoneal injection method, intragastric gavage of riluzole significantly improved the fertility of oligospermic mice induced by the BSF, resulting in a fertility rate of 66.67% (Figure 2B). Semen analysis revealed significant improvements in various sperm quality parameters, such as count, viability, motility, and abnormality rate (Figure 2C-G). Furthermore, riluzole therapy effectively increased testicular and epididymis weight compared to BSF-induced group (testes:102.10 ± 15.59 mg vs. 42.55 ± 2.89 mg; epididymis: 37.28 ± 2.46 mg vs. 26.30 ± 0.79 mg) (Figure 2H-I). Histological examination of testicular sections demonstrated that the intragastric gavage administration of riluzole effectively rescued testicular atrophy and countered the reduction in germ cell numbers (Figure 2J-K). It also improved the Johnsen score of the seminiferous epithelium and increased the number of mature sperm in the epididymis (Figure 2L-N). These findings confirmed that the switch to intragastric gavage administration of riluzole not only eliminated the risk of abdominal organ damage and improved patient compliance but also maintained the therapeutic effects observed with the previous intraperitoneal injection method.

**Figure 2.**
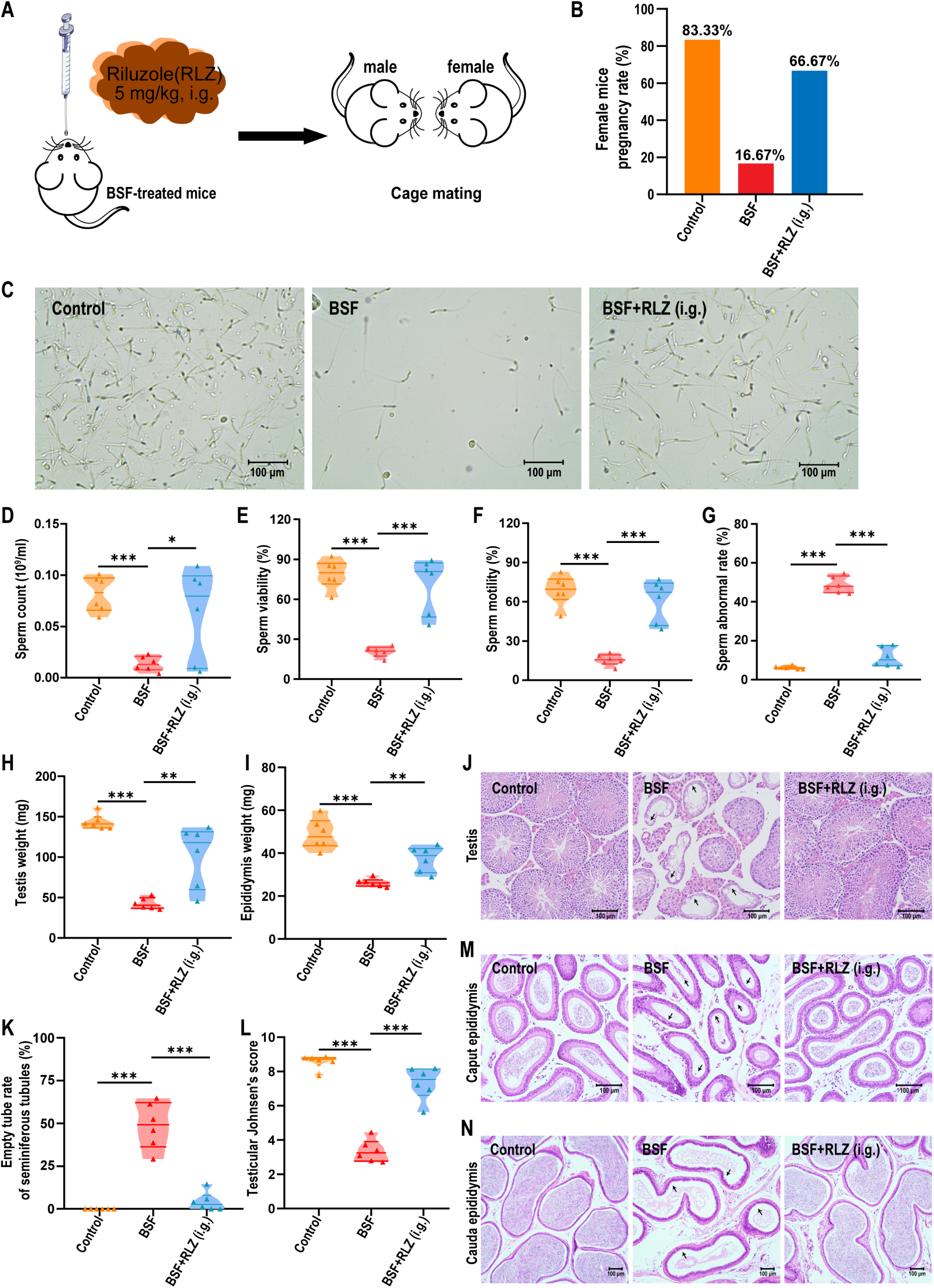
Intragastric gavage administration of riluzole rescued fertility and sperm quality in oligospermic mice. (**A**) Schematic diagram of intragastric gavage administration to BSF-treated mice. (**B**) Fertility rate of mice with oligospermia induced by BSF after 8 weeks of riluzole treatment via intragastric gavage (5 mg/kg·bw) (n = 6). (**C**) Representative micrographs of sperms. (**D-G**) Count, viability, motility and abnormal rate of sperms in the cauda epididymis of oligospermic mice (n = 6). The data were analyzed for more sperm counts over 1000 with CASA. (**H-I**) Testis and epididymis weight in mice (n =6). (**J**) Representative histopathology of the testes. Black arrow presented low numbers of germ cells in the seminiferous tubule lumen. Scale bar, 100 μm. (**K-L**) Empty tube rate and testicular Johnsen’s score of seminiferous tubules in oligospermic mice after riluzole therapy (n=6). (**M-N**) Representative histopathology of the caput epididymis and cauda epididymis. Black arrow presented lower density of sperms in the epididymis. Scale bar, 100 μm. Values are expressed as mean ± SEM. One-way analysis of variance (ANOVA) was used to analyze statistical differences; \*\*\**P* < 0.001, \*\**P* < 0.01, \**P* <0.05 compared to BSF.

### Riluzole administration enhanced spermatogonia differentiation and regulated blood-testis barrier function in BSF-treated mice

To determine the role of riluzole in the differentiation of spermatogonia, we examined the expressions of GFRα1, PCNA, SYCP3, and TNP-1, key markers involved in spermatogonia proliferation and differentiation. Our findings revealed that a significant decrease in the expression levels of GFRα1 (which is associated with the self-renewal of SSCs (Shamhari et al., 2023)), PCNA (a marker for proliferating cell nuclear antigen (Fu et al., 2019)), SYCP3 (involved in meiosis (Miyamoto et al., 2003)), and TNP-1 (specific to spermatids (Della-Maria, Gerard, Franck, & Gerard, 2002)) after BSF treatment. However, treatment with riluzole, whether administered via intraperitoneal injection or intragastric gavage, significantly increased the expression of these marks (Figure 3A-D). These findings suggested that riluzole promoted the proliferation, differentiation, and maturation of spermatogonia during spermatogenesis. We investigated the impact of riluzole treatment on the tight junctions of the blood-testis barrier (BTB) in BSF-treated mice by focusing on SCs, which are responsible for forming the BTB and protecting spermatogonia proliferation and differentiation as well as sperm regeneration. To assess BTB permeability, we utilized sulfo-NHS-LC-biotin, a water-soluble and membrane-impermeable biotinylation reagent. In normal mouse testes, sulfo-NHS-LC-biotin did not enter the seminiferous tubules. However, after BSF treatment, the presence of sulfo-NHS LC-biotin within the tubules indicated disruption of BTB tight junctions. Remarkably, riluzole treatment restored the integrity of the BTB, thereby preventing the entry of sulfo-NHS LC-biotin into the seminiferous tubules (Figure 3E).

**Figure 3.**
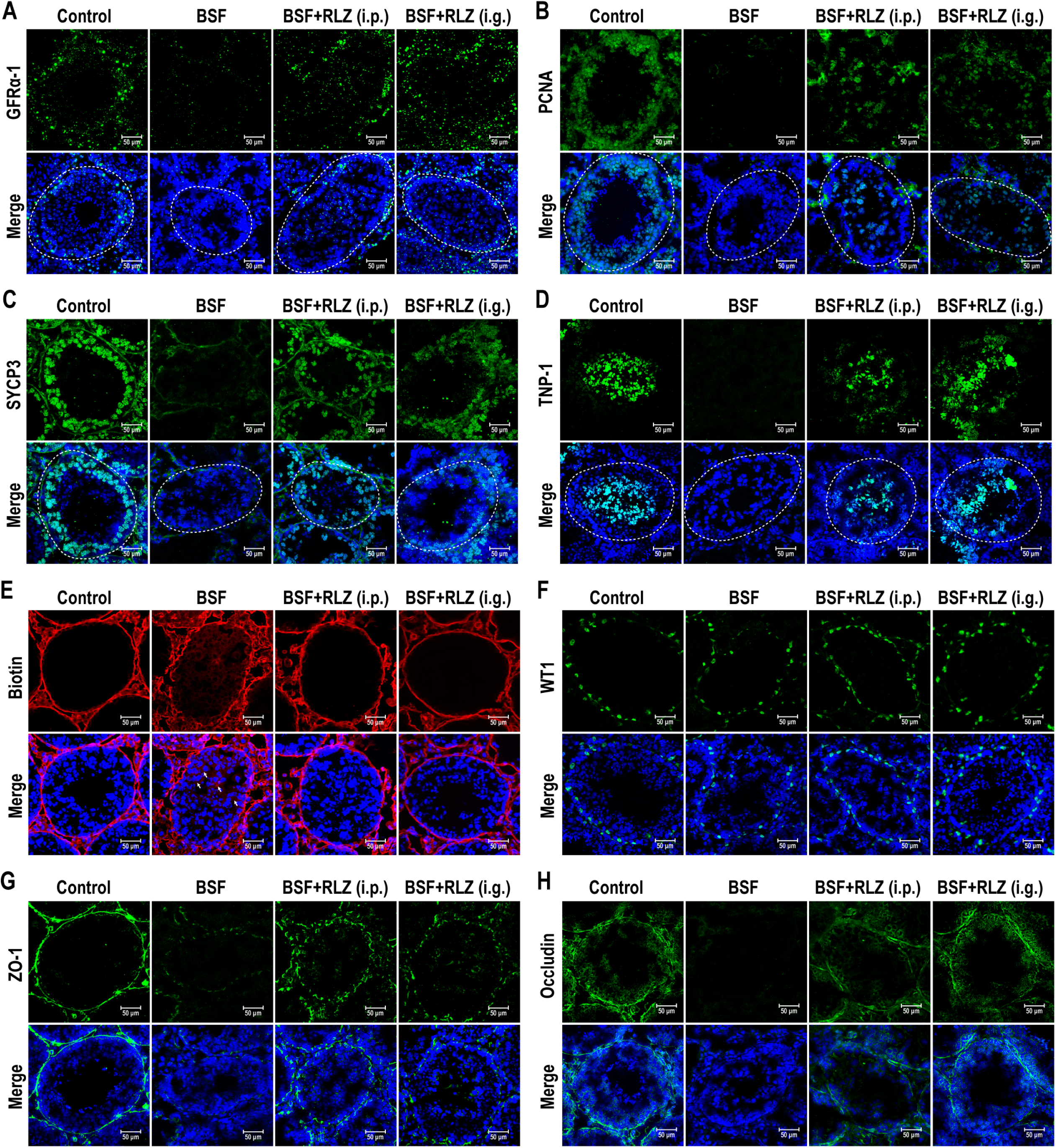
Effects of riluzole on proliferation and differentiation of spermatogonia and BTB integrity. (**A-D**) Immunofluorescence staining showed the expressions of spermatogenesis-related proteins (GFRα-1, PCNA, SYCP3, and TNP-1) in the testis of BSF induced oligospermic mice after riluzole treatment. Nuclei were stained with DAPI (blue). Scale bar, 50 μm. (**E**) Assessment of BTB integrity using the sulfo-NHS-LC-biotin assay. The nuclei were stained with DAPI (blue). White arrows indicated the presence of biotin in the lumen of seminiferous tubules, indicating BTB disruption. Scale bars, 50 μm. (**F-H**) Immunofluorescence staining showed the expressions of WT1, a marker for Sertoli cells, and proteins related to the blood-testis barrier (ZO-1 and Occludin) in the testes. Nuclei were stained with DAPI (blue). Scale bar, 50 μm.

Furthermore, we evaluated the expression of WT1 (SCs marker (Y. Wen et al., 2021)) and the tight junction-related proteins ZO-1 and Occludin (Gonzalez-Mariscal et al., 2000). Our findings revealed a significant decrease in the expression of ZO-1 and Occludin after BSF treatment, while WT1 expression remained unaffected. This suggested that BSF primarily targeted the disrupting of BTB integrity. In contrast, riluzole treatment increased the expression of ZO-1 and Occludin, suggesting its potential to restore BTB damage by upregulating the expression of tight junction-related proteins (Figure 3F-H). In conclusion, these findings indicated that riluzole enhanced sperm regeneration by restoring the integrity in the BTB.

### Transcriptomics analysis revealed the regulatory mechanism of riluzole on tight junctions in BSF-treated mice

Transcriptomic analysis was performed to illustrate how riluzole regulated the testicular gene expression patterns. The Pearson correlation coefficient square (R^2^) indicated the clustering relationship among the normal mice (Control group), BSF-induced oligospermic mice (BSF group), and BSF-induced oligospermic mice treated with riluzole (BSF+RLZ group). The BSF group was distinct from the Control group, while the riluzole treatment group clusters with the Control group (Figure 4A). The global gene expression patterns of riluzole treatment group resemble those of the Control group (Figure 4B). In comparison to the BSF group, the Control group exhibited 1933 up-regulated genes, and 3843 down-regulated genes, while the riluzole treatment group exhibited 1781 up-regulated genes and 3108 down-regulated genes (Figure 4C-D). A venn diagram showed that there were 4105 common differentially expressed genes (DEGs) among the two comparison groups: the Control vs BSF group and riluzole treatment vs BSF group (Figure 4E). Gene Ontology (GO) analysis, including biological process (BP), cellular component (CC) and molecular function (MF) and Kyoto Encyclopedia of Genes and Genomes (KEGG) pathway enrichment analysis were conducted on the 4105 common DEGs for the identification of key regulatory genes affected by riluzole. The riluzole treatment leaded to differential gene expression associated with various functions of the seminiferous cord (SC), including extracellular matrix binding, cell migration, and cell adhesion. Additionally, it affected specific cellular components, such as acrosomal vesicles, sperm flagellum, sperm plasma membrane and sperm head (Figure 4F). KEGG pathway enrichment analysis highlights focal adhesion, axon guidance, peroxisome, apoptosis, regulation of actin cytoskeleton, tight junction, drug metabolism, calcium signaling pathway, chemokine signaling pathway and gap junction as the primary enriched pathways closely associated with SCs function and blood-testis barrier (Figure 4G).

**Figure 4.**
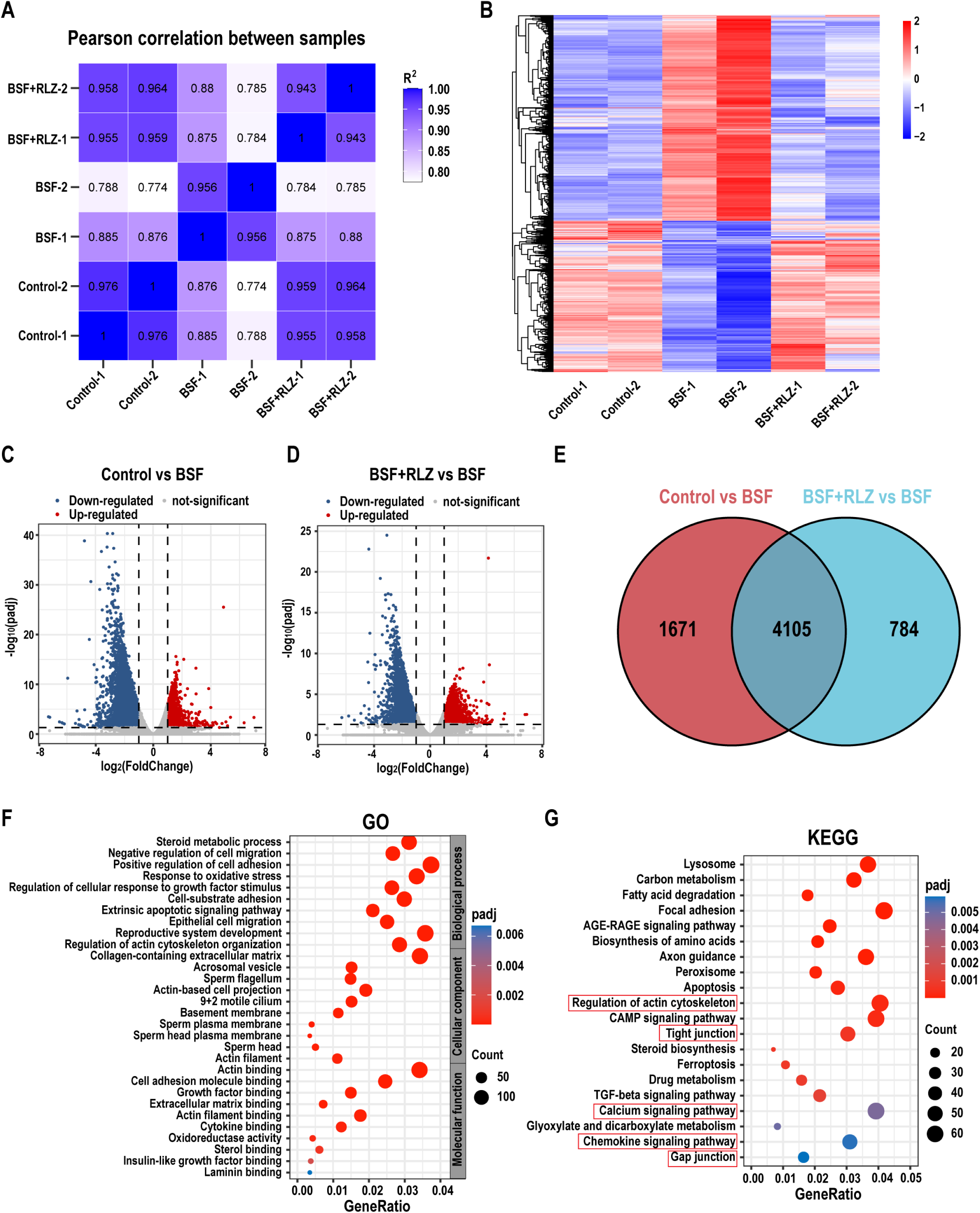
Changes in the transcriptome profile of testis after riluzole treatment. (**A**) The Pearson correlation coefficient was exhibited as coefficient values to determine the correlation of the transcriptional profiles between each sample. (**B**) Cluster analysis of differentially expressed genes (DEGs) expression changes in testis between each group. The color bar indicates gene expression on a log_2_ scale. Red represented up-regulated and blue represented down-regulated gene expression. (**C-D**) Volcano plot of the DEGs of testes between each group. The criterion for screening DEGs had adjusted padj < 0.05 and | log_2_FoldChange | ≥ 1. Up-regulated genes were represented by red dots, down-regulated genes were represented by blue dots. Gray dots indicate genes with no difference in expression. (**E**) Venn diagram analysis of the differential gene. (**F-G**) GO and KEGG pathway enrichment analysis of the DEGs. The abscissa in the figure is the ratio of the number of differential genes annotated to the GO Term or KEGG pathway to the total number of differential genes, and the ordinate is the GO Term or KEGG pathway. The size of the point represented the number of genes annotated to the GO Term or KEGG pathway. The color of the dot indicated the enrichment degree (padj) of the pathway. Control group: normal mice; BSF group: BSF-induced oligospermic mice; BSF+RLZ group: BSF-induced oligospermic mice treated with riluzole.

### Riluzole enhanced the integrity of the BTB by improving the functioning of Sertoli cells

To investigate the potential role of riluzole in restoring BTB integrity, we examined its effects on SCs proliferation, paracrine function, migration ability, and ability to form tight junctions. Firstly, we assessed the impact of riluzole on SCs proliferation. Our results showed that concentrations of riluzole below 20 μM had no significant effect on cell viability (Figure 5A). Next, we investigated the influence of riluzole on the paracrine function of SCs by treating them with 10 μM riluzole for 24 h. The expression levels of key genes involved in spermatogenesis, including *Gdnf*, *Bmp4*, *Scf*, *Cxcl12*, *Inhibin B*, and *Fgf2,* were examined. Results revealed that riluzole significantly upregulated the expression of these genes (Figure 5B). We also investigated the effect of riluzole on SCs migration, as this ability is essential for regulating BTB damage repair. We compared the migration ability of SCs with and without riluzole treatment. BSF inhibited the migration ability of SCs, while riluzole restored it, indicating that riluzole could promote the migration of SCs (Figure 5C-D). Furthermore, we examined the ability of riluzole to regulate the formation of tight junctions in the blood-testis barrier. The results suggested that riluzole significantly upregulated the expression of blood-testis barrier-related proteins ZO-1 and Cx43 (Figure 5E-I). To assess the permeability of SCs monolayers, the Transepithelial Electrical Resistance (TER) across these layers was measured. BSF markedly lowered the TER value, which suggested a disruption in the integrity of the Sertoli cell barrier. In contrast, treatment with riluzole significantly increased the TER value, implying a restoration of barrier integrity (Figure 5J).

**Figure 5.**
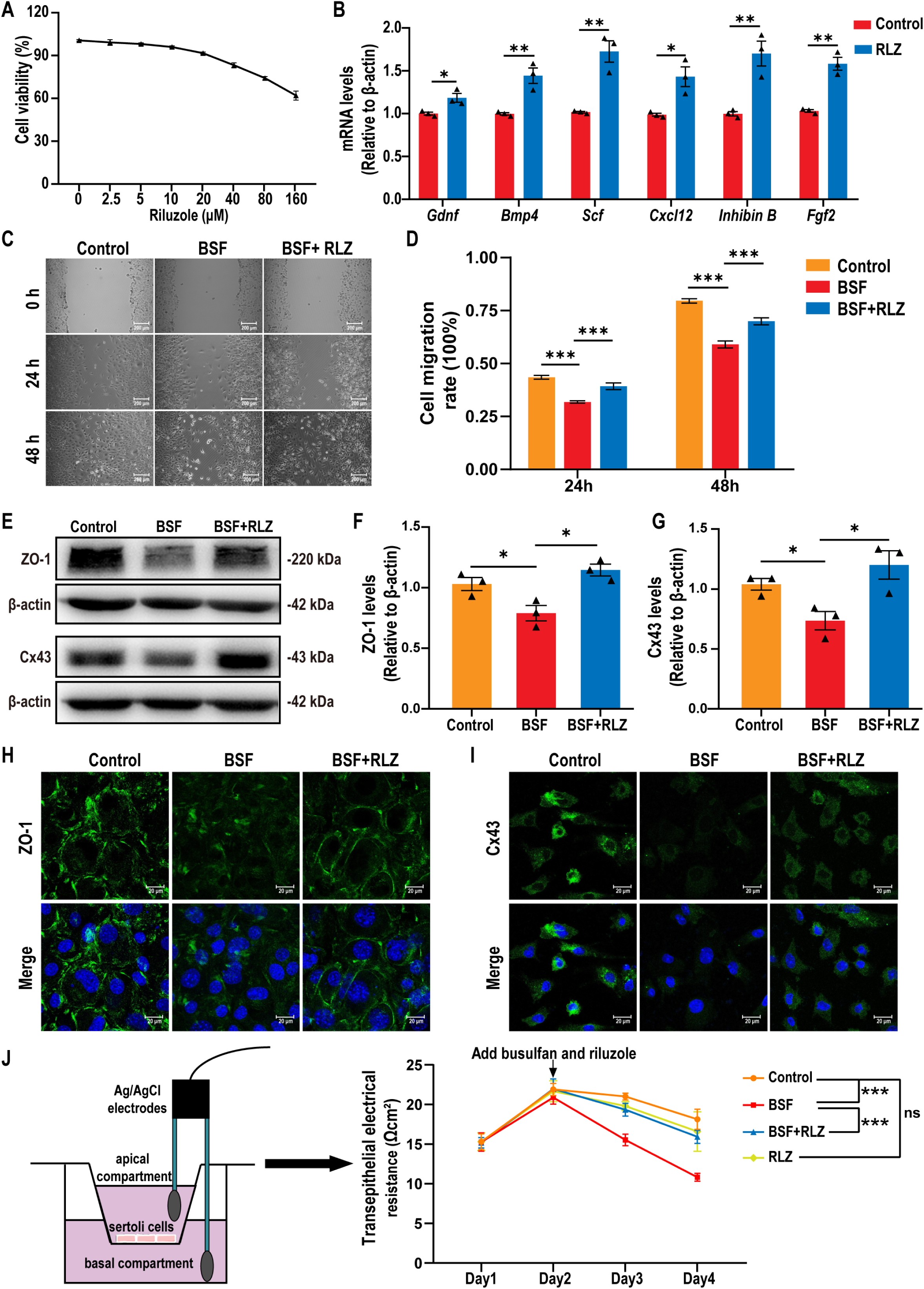
Effects of riluzole on Sertoli cells (SCs) functionality in vitro. (**A**) The cell viability of SCs treated with different concentrations of riluzole. (**B**) Expression levels of SC-related secretory factors (*Gdnf*, *Bmp4*, *Scf*, *Cxcl12*, *Inhibin B*, and *Fgf2*) in SCs treated with 10 μM riluzole for 24 h, measured by qPCR with β-actin as the internal reference. (**C**) Scratch assay showing the effect of 10 μM riluzole and/or 200 μM BSF on SCs migration at 24 and 48 h. Representative images captured under a microscope. Scale bar, 200 μm. (**D**) Cell migration rate calculated based on average scratch distance using Image J analysis after 24 and 48 h of treatment. (**E-G**) Western blot analysis of ZO-1 and Cx43 expression in SCs treated with 10 μM riluzole and/or 200 μM BSF, with β-actin as the loading control. (**H-I**) Immunofluorescent staining for ZO-1 and Cx43 in SCs treated with 10 μM riluzole and/or 200 μM BSF. Nuclei were stained with DAPI (blue). Scale bar, 20 μm. (**J**) Transepithelial electrical resistance (TER) measurement to assess Sertoli cell TJ-permeability barrier function. All data were obtained from three independent experiments and presented as mean ± SEM. Statistical significance was determined by two-tailed Student’s t test (B) or one-way ANOVA (D, F-G, J); \*\*\**P* < 0.001, \*\**P* < 0.01, \**P* <0.05.

Overall, these findings suggested that riluzole had the potential to restore BTB integrity by promoting Sertoli cell proliferation, enhancing the paracrine function,restoring migration ability, and promoting the formation of tight junctions.

### Riluzole restored BTB function by activate the transient receptor potential canonical subfamily member 5 (TRPC5)

The transcriptome profile analysis revealed that the calcium signaling pathway played a role in the regulation of BTB function by riluzole. Additionally, it has been known that the regulation of intracellular calcium is critical for the normal function of tight junctions (Stuart et al., 1994). In light of this, we specifically examined the expression changes of differentially expressed genes (DEGs) in the calcium signaling pathway. Out of the 62 DEGs identified, 12 genes (*Grin2b*, *Grin3b*, *Smim6*, *Calml4*, *Hgf*, *Ntrk3*, *Plcz1*, *Pln*, *Pde1a*, *Calml3*, *Camk4*, and *P2rx3*) were up-regulated in the riluzole treatment group compared to the BSF group. Conversely, 50 genes were down-regulated (Figure 6A). To explore the potential interactions among these DEGs, we constructed a protein-protein interaction (PPI) network using the STRING database. The PPI network analysis revealed a single cluster, suggesting that these genes may be regulated by common mechanisms (Figure 6B).

**Figure 6.**
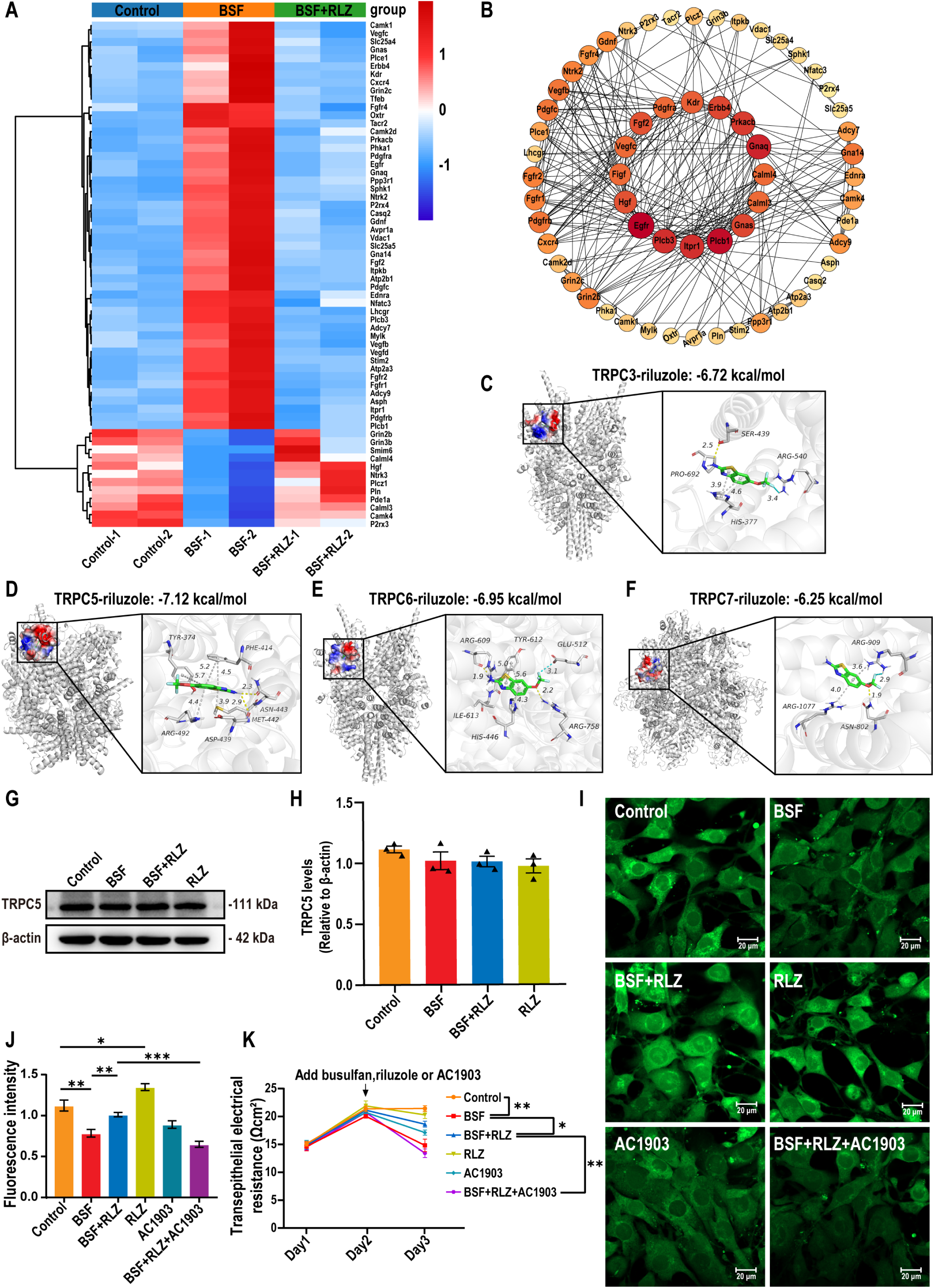
Riluzole activated TRPC5 and regulated intracellular calcium levels. (**A**) Heat map showed the changes in gene expression in calcium signaling pathway. The red represented up-regulated and the blue represented down-regulated gene expressions. (**B**) Protein-protein interaction (PPI) analysis of the DEGs involved in calcium signaling pathway from testes. Network visualized with Cytoscape. (**C-F**) The binding mode of TRPC3/5/6/7 with riluzole. (c) TRPC3-riluzole; (d) TRPC5-riluzole; (e) TRPC6-riluzole; (f) TRPC7-riluzole. On the left was the electrostatic surface of complex, and on the right was the detail binding mode of complex. Yellow dash, gray dash, blue dash showed the hydrogen bond, π-stacking interaction, and halogen bond, respectively. (**G-H**) Western blot analysis of Sertoli cells (SCs) treated with 200 μM BSF and/or 10 μM riluzole to detect TRPC5 expression. β-actin was used as the internal reference. (**I-J**) Changes in intracellular calcium concentration. (**K**) TER to assess changes in the function of the SCs tight junction-permeability barrier. All data were obtained from three independent experiments and presented as mean ± SEM. One-way analysis of variance (ANOVA) was used to analyze statistical differences; \*\*\**P* < 0.001, \*\**P* < 0.01, \**P* <0.05.

To further clarify the molecular mechanism of riluzole in regulating calcium signaling pathways, we conducted molecular docking to predict the interaction between riluzole and TRPC (Transient Receptor Potential Canonical) proteins. TRPC is a nonselective calcium-permeable cation channel responsible for the entry of calcium ions (Ca^2+^) into cells (Numaga-Tomita & Nishida, 2020). The TRPC subtypes analyzed in our study included TRPC3, TRPC5, TRPC6, and TRPC7. Docking models of riluzole with these subtypes revealed binding energies of −6.72 (TRPC3), −7.12 (TRPC5), −6.95 (TRPC6), and −6.25 (TRPC7) kcal/mol, respectively. Importantly, the binding energy between riluzole and TRPC5 was found to be higher than that with TRPC3, TRPC6, and TRPC7, indicating a stronger affinity between riluzole and TRPC5. And riluzole comprises hydrophobic benzene rings that form robust hydrophobic interactions with TRPC5 through active site amino acids, such as PHE-414. The benzene rings of riluzole also exhibit π-stacking interactions with PHE-414, TYR-374, and MET-442. Furthermore, riluzole forms strong hydrogen bond interactions with the ASP-439 and ASN-443 sites of TRPC5 (Figure 6C-F).

To further investigate the role of TRPC5 in the regulation of BTB tight junctions by riluzole, the expression of TRPC5 in SCs was examined. The results confirmed the presence of TRPC5 in SCs; however, no significant changes in TRPC5 expression were observed following treatment with riluzole or BSF (Figure 6G-H). Subsequently, the intracellular calcium levels in SCs were assessed. Treatment with BSF caused a decrease in intracellular calcium ion concentration, while riluzole treatment induced an increase in intracellular calcium ions, effectively counteracting the inhibitory effect of BSF on calcium ion levels. Importantly, the effect of riluzole on intracellular calcium levels was blocked by AC1903, a specific and selective inhibitor of TRPC5. This strongly suggested that riluzole modulates calcium signaling through the activation of TRPC5 (Figure 6I-J). Additionally, the effect of riluzole on the integrity of tight junctions was evaluated using transepithelial electrical resistance (TER) assays. The results showed that riluzole significantly increased the TER value, indicating an enhancement of tight junction function. Notably, the increase in TER induced by riluzole was reversed by AC1903 (Figure 6K). Thus, these findings suggest that riluzole promotes the integrity and function of tight junctions by activating TRPC5.

### Riluzole had no significant adverse effects on male mice reproduction health

To assess the potential adverse impact of riluzole on the reproductive ability of male mice, riluzole was administered through intragastric gavage at a daily dose of 5 mg/kg for a duration of 90 days (Figure 7A). The results indicated that there were no significant differences in body and reproductive organ weights (testis and epididymis) when comparing the exposed mice to the control group (Figure 7B-D). Additionally, the analysis of sperm parameters, including count, viability, motility, and abnormality ratio, revealed no significant damage to sperm quality in male mice exposed to riluzole (Figure 7E-H). Moreover, the fertility rate and number of offspring in the exposed male mice were comparable to those in the control group (Figure 7I-J). Histological examination showed no observable damage to the testis and epididymis of male mice treated with riluzole (Figure 7K-L). In summary, these findings provide evidence that the long-term administration of riluzole does not have any noticeable toxic effects on the reproductive capacity of male mice.

**Figure 7.**
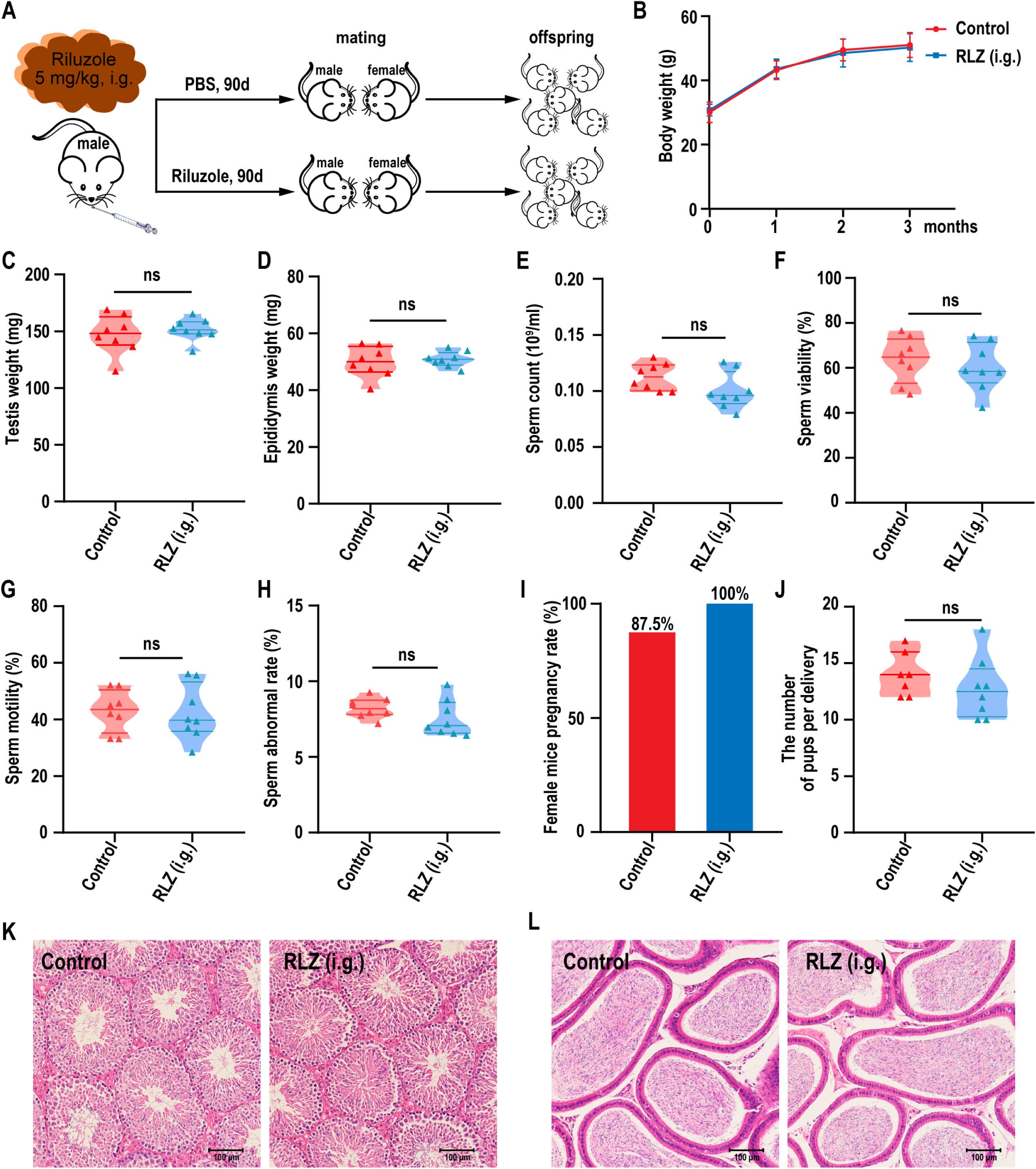
Assessment of the reproductive ability of male mice exposed riluzole for 90 days via intragastric gavage. (**A**) Illustration of experimental scheme. (**B**) Body weight of male mice after riluzole exposed through intragastric gavage at a daily dose of 5 mg/kg for a duration of 90 days (n = 8). (**C-D**) Testis and epididymis weight in male mice exposed to riluzole (n = 8). (**E-H**) Sperm count, viability, motility and abnormal rate in male mice exposed to riluzole (n = 8). The data were analyzed for more sperm counts over 1000 with CASA. (**I**) Fertility rate in male mice exposed to riluzole (n = 8). (**J**) Number of offspring of male mice exposed to riluzole. (**K-L**) Representative histopathology of the testis and epididymis of male mice exposed to riluzole. Scale bar, 100 μm. Values were presented as mean ± SEM. Two-tailed Student’s t test was used to analyze statistical differences; ns: no significance.

## Discussion

In our study, we have identified that riluzole has the ability to regulate SCs functions, leading to improved spermatogenesis and restored fertility in oligospermic mice induced by BSF. Moreover, our research indicated that riluzole achieved these effects by modulating intracellular calcium levels in SCs. We identified that riluzole binds to TRPC5, a calcium-permeable channel involved in the regulation of intracellular calcium (Richter, Schaefer, & Hill, 2014). By binding to TRPC5, riluzole can effectively modulate calcium levels in SCs, leading to the observed improvements in spermatogenesis and fertility (Figure 8). These findings highlight the potential of riluzole as a therapeutic intervention for chemotherapy-induced male reproductive damage.

**Figure 8.**
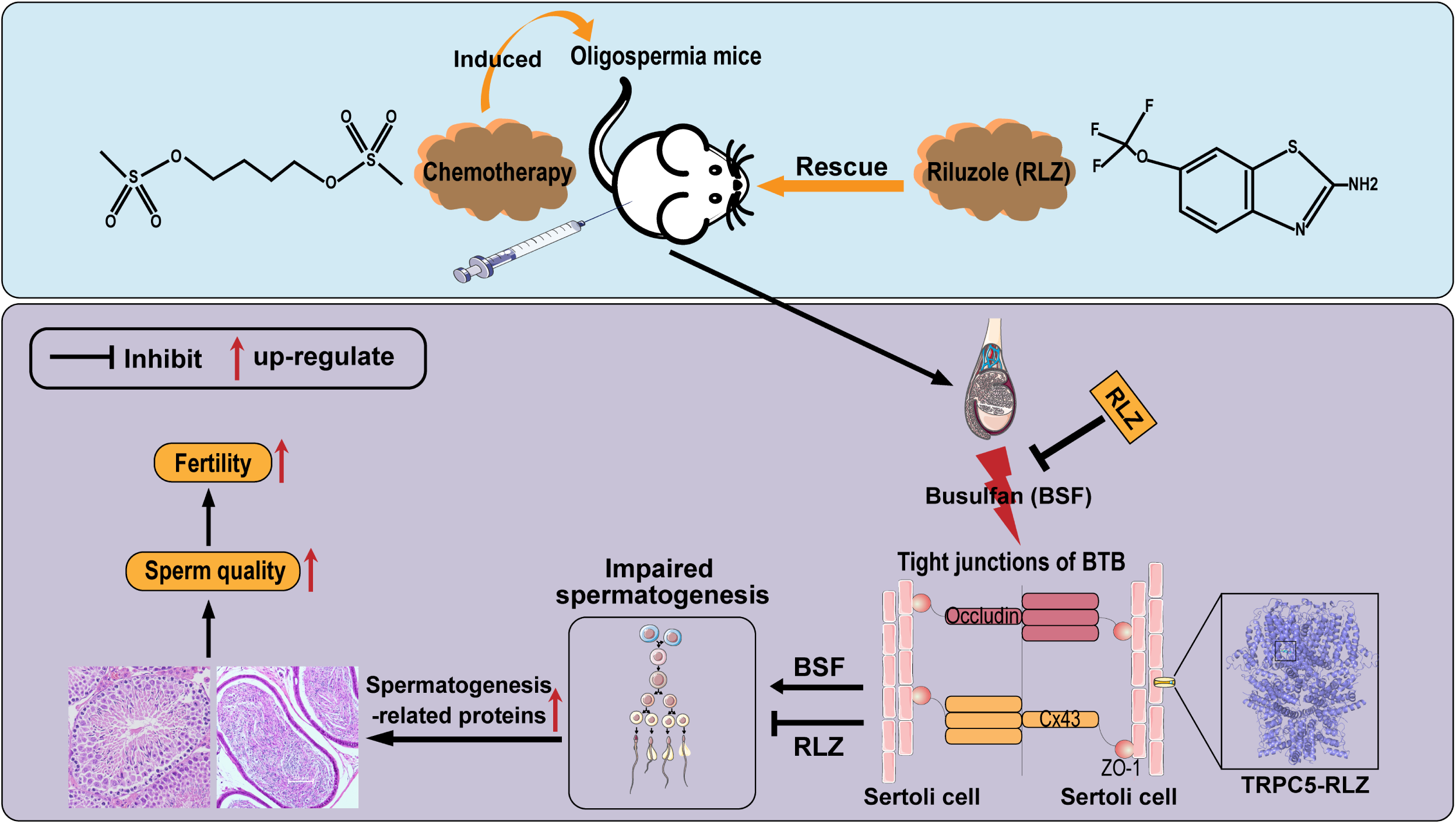
Schematic diagram of the potential mechanism of RLZ against BSF-induced reproductive injury. Riluzole modulates the blood-testis barrier (BTB), enhancing spermatogenesis and restoring fertility in mice with busulfan (BSF)-induced oligospermia through its interaction with TRPC5.

Riluzole is a well-studied drug primarily used for the treatment of amyotrophic lateral sclerosis (ALS) and has been shown to possess neuroprotective properties (Jaiswal, 2019; Miller, Mitchell, & Moore, 2012; Wokke, 1996). Its mechanism of action involved blocking ion channels and inhibiting glutamate activity, leading to the regulation of glutamate-dependent signaling (Doble, 1996; Fumagalli, Funicello, Rauen, Gobbi, & Mennini, 2008; Seol et al., 2016). This mechanism has been implicated in various neurological conditions, including ischemic brain injury, Parkinson’s disease, Alzheimer’s disease, and Huntington’s disease (Bensimon et al., 2009; Liu & Wang, 2018; Squitieri, Ciammola, Colonnese, & Ciarmiello, 2008; Vallée, Vallée, Guillevin, & Lecarpentier, 2020; Verma, Arora, Javed, Akhtar, & Samim, 2016). Furthermore, riluzole has demonstrated therapeutic efficacy in epilepsy and certain psychiatric disorders such as bipolar depression and anxiety (Diao et al., 2013; Lapidus, Soleimani, & Murrough, 2013; Mathew et al., 2005; Park et al., 2017). It modulated the Wnt/β-catenin signaling pathway by inhibiting glycogen synthase kinase-3β (GSK-3β) opens up potential applications in cell self-renewal, embryonic development, and organogenesis (Biechele et al., 2010; Y. Yang, Li, et al., 2022). Despite the extensive research on riluzole, its impact on spermatogenesis has not been extensively explored. In our previous study, we found that riluzole had the ability to induce the transformation of mouse embryonic fibroblasts into Sertoli-like cells (Y. Yang, Li, et al., 2022). These Sertoli-like cells showed increased expression of key cytokines (*Gdnf*, *Bmp4*, *Scf*, *Cxcl12*, *Inhibin B*, and *Fgf2*) involved in the regulation of spermatogonial stem cell proliferation and differentiation. For example, GDNF was found to promote the self-renewal of stem cells, while BMP4 acted on their differentiation (Hu et al., 2004; Meng et al., 2000). SCF regulated the fate of type A spermatogonia, and CXCL12 was involved in the migration of stem cells (Feng, Ravindranath, & Dym, 2000; Q. E. Yang, Kim, Kaucher, Oatley, & Oatley, 2013). Based on these observations, we hypothesized that riluzole might improve spermatogenesis by enhancing the function of SCs. To test this hypothesis, we conducted an in vitro experiment using testicular tissue cultured in the presence of BSF. Treatment with riluzole for 7 days showed potential in restoring the reduction in the number of germ cells in the seminiferous tubules (data are shown in Supplementary file 2). This finding suggested that riluzole could potentially be used as a therapeutic option for oligospermia, a condition characterized by low sperm count (McLachlan, 2013). Therefore, we aim to further investigate the relationship between riluzole and spermatogenesis, potentially uncovering new applications for this drug in the field of male infertility.

To investigate whether riluzole promotes spermatogenesis through regulating the HPG axis, we examined its effect on sex hormones. We found that there was no significant difference in sex hormone levels compared to the control group (data are shown in Supplementary file 3). This suggested that riluzole did not directly activate the HPG axis or increase sex hormone levels to promote sperm regeneration in BSF-induced oligospermic mice. Following this, we examined the effect of riluzole on SCs. Our study revealed that riluzole enhanced the integrity of the BTB by improving the functioning of SCs. Transcriptomic analysis showed that calcium signaling pathways play a significant role in this process. Previous research has shown that riluzole binds near a calcium ion and can activate TRPC5 (Y. Yang, Wei, & Chen, 2022). Our research validates that riluzole specifically binds to TRPC5, leading to the activation of calcium signaling pathways. This, in turn, enhances the expression of tight junction proteins, particularly ZO-1 and Cx43. These proteins play a key role in interacting with the actin cytoskeleton and forming linkages with various junction types within the BTB (Cheng & Mruk, 2012; Segretain et al., 2004). This understanding of the molecular mechanism behind riluzole’s effects on the BTB can pave the way for further research and the development of targeted interventions for male infertility caused by chemotherapy. More importantly, our toxicity evaluation results have shown that long-term administration of riluzole for 90 days has no significant impact on the reproductive ability of mice. This suggests that riluzole is safe for use in the reproductive system.

The riluzole is already an approved drug for ALS. Its safety profile is well-established, and it has been used in clinical practice for several years (Brooks, Bettica, & Cazzaniga, 2019; Pongratz, Neundörfer, & Fischer, 2000; Salardini et al., 2016). This provides an advantage in terms of streamlined evaluation of safety and tolerability in repurposing riluzole for oligospermia treatment. The potential to expedite the approval process and speed up the availability of an effective treatment for oligospermia is significant when considering the urgency of addressing male infertility. In addition to safety and timeliness, repurposing riluzole could also offer cost and resource savings. Developing novel drugs from scratch is a lengthy and expensive process. By utilizing existing drugs like riluzole, we can leverage the extensive research and development already conducted, reducing costs and saving valuable resources. This approach can potentially make the treatment more affordable and accessible to a larger population.

However, more studies are required to elucidate the mechanisms underlying these beneficial effects and to determine the long-term safety and efficacy of riluzole in male reproductive health. Clinical trials involving larger sample sizes and diverse populations are necessary to validate these initial findings and assess the generalizability of the results. Moreover, it is important to consider potential side effects or interactions of riluzole with other medications commonly used in chemotherapy regimens. Therefore, careful evaluation and monitoring would be essential before considering riluzole as a standard treatment option for chemotherapy-induced oligospermia.

In conclusion, our findings provide insight into the underlying mechanisms through which riluzole exerts positive effects on male reproductive outcomes. This knowledge has the potential to guide future research and clinical trials in exploring the full therapeutic potential of riluzole for treating oligospermia and other related male reproductive conditions. Repurposing old drug riluzole for new uses could expedite the development of new treatments for fertility-related complications caused by cancer therapy and reduce the risks and costs associated with developing completely new drugs.

## Materials and methods

### Animals

Healthy male kunming mice aged at 4 weeks were purchased from the Experimental Animal Center of Guangdong Province (Guangzhou, China). Before the experiments, all of the animals were acclimatized at least 7 days under a 12 h light/dark cycle with ad libitum access to food and water in a controlled temperature (24 ± 2 °C) with relative humidity (50%-60%). Except where otherwise stated, singly caged during experimentation to prevent them from fighting. All animal experiments were approved by the Institutional Animal Care and Use Committee (IACUC) of Jinan University (IACUC Approval: 20200327-68). The National Institutes of Health Guide for the Care and Use of Laboratory Animals was adhered to during the implementation of all animal experimental.

### Intraperitoneal and intragastric gavage administration of riluzole

After one week of acclimatization, the mice accepted a single intraperitoneal injection of busulfan (BSF, Cat# HY-B0245, MedChemExpress, New Jersey, USA) at the doses of 30 mg/kg. After a week, mice were randomly divided into groups (six mice in each group) and treated with riluzole (3 mg/kg·bw for intraperitoneal injection, and 5 mg/kg·bw for intragastric gavage administration) once a day for 1 week. Riluzole (Cat# S1614, Selleckchem, Houston, Texas, USA) was dissolved in DMSO to create an experimental dosage stock solution. The riluzole stock solution was further dissolved with 40% PEG300 (Cat# HY-Y0873, MedChemExpress), 5% Tween 80 (# HY-Y1891, MedChemExpress) and 45% PBS for use in intraperitoneal injection. For intragastric gavage administration, riluzole oral solution contains 2% riluzole stock solution and 98% PBS. Also, the Control group was treated with the corresponding solvent and raised under the same conditions to serve as the basic control (See Supplementary file 4). 6 weeks later, all mice were weighed before being euthanized, followed by the collection of blood and tissue samples. All of the mice survived until the finalization of the drug administration. No criteria were set for including or excluding animals and there was no exclusion of mice in this study.

### Mating with female mice test

After 5 weeks of riluzole administration, male mice were mated with adult estrous kunming females for 7 days. Females of pairs breeding under normal circumstances were then monitored for the presence of vaginal plugs and pregnancies.

### Semen analysis

Sperm masses were obtained from the epididymis cauda using a previously described protocol (Luo et al., 2018). Briefly, the cauda epididymis was put into a 1.5 mL microcentrifuge tube containing 1 mL PBS and simultaneously cut into small pieces for an entire sperm releasing at a sustaining temperature of 37 ℃ for 15 min. The sperm suspension was then loaded in a sperm counting chamber and analyzed by a computer-aided semen analysis system (CASA) (Malang ML-608JZII, Nanning, China). Images of 30-40 fields of the chamber (about 1000 sperms) each mouse were recorded randomly and the index including sperm count, viability, motility, abnormality, and other motion parameters was examined for a comprehensive assessment to sperm masses.

### Histopathology of the testicles and epididymis

All fixed testes and epididymis were embedded in paraffin after dehydrating in a graded series of ethanol and sectioned into slides with a thickness of 5 μm. Then the slides were stained by hematoxylin and eosin (H&E). After final dehydrated through a graded series of alcohol and soaked in xylene for 10 min, slides were mounted using neutral gum. H&E staining slides were visualized under a light microscope (Nikon, Tokyo, Japan). To evaluate the changes of seminiferous epithelium, the spermatogenic tubules score was conducted according to the standard of testicular modified Johnsen score as described previously (Ito et al., 2021). At least 50 spermatogenic tubules were analyzed in each mouse.

### Immunofluorescence of tissue

For immunofluorescence experiments, testes were embedded in Optimum Cutting Temperature (OCT) compound and cryosectioned at a thickness of 6 μm. The sections were permeabilized by incubation with 1% Triton X-100 in PBS for 30 min at room temperature. Non-specific adhesion sites were blocked with blocking buffer (Cat# P0102, Beyotime, Shanghai, China) for 60 min at room temperature. Then the sections were incubated with primary antibodies at 4 ℃ overnight, followed by secondary antibody conjugated to Alexa Fluor 488 (Cat# ab150077, Abcam) for 1 h at room temperature. Nuclei were stained with DAPI (Cat# AR1177, Bosterbio). Stained samples were then visualized, and images were captured using a LSM900 confocal microscope (Zeiss, Germany). Details of the primary antibody were listed in Supplementary file 5.

### Sertoli cells (SCs) isolation and treatments

Sertoli cells (SCs) were isolated from the testes of 7-day-old male mice. Briefly, the testes denuded of tunica albuginea were incubated with 1 mg/mL collagenase type IV in DMEM for 7 min at 37 °C water bath and then centrifuged at 100 × g for 2 min to eliminate interstitial cells. The centrifuged seminiferous tubules were further digested with 0.25% trypsin-EDTA for 10 min at 37 °C in a water bath and then filtered through a 40 µm cell strainer. The cells in the filtrate were collected by centrifugation (250 × g, 5 min) and resuspended in DMEM with 10% fetal bovine serum. Next, the cell suspensions containing primary SCs and spermatogonia were cultured in cell culture dish at 37 °C with 5% CO_2_. Four hours later, the culture supernatant was collected to remove spermatogonia, and the adherent cells were pure SCs we need. After 48 h of routine culture, SCs were seeded into 6-well plates (5 × 10^5^ cells/well), allowed to adhere 24 h to crosslinking approximately 90%, and media was replaced with serum free media to deprivation 2 h before riluzole and BSF treatment. Then, cells were treated with 200 μM BSF and / or 10 μM riluzole. After 24 h of treatment, cells were collected for real-time PCR analysis and western blot analyses.

### Quantitative RT-PCR (qRT-PCR)

Total RNA was extracted from Sertoli cells using TRIzol reagent (Cat# 15596018, Invitrogen, Carlsbad, CA, USA). One microgram of total RNA was reverse transcribed into cDNA using PrimeScript™ RT Master Mix (Cat# RR036A, TaKaRa, Japan). qPCR was conducted with ChamQ SYBR qPCR Master Mix (Cat# Q311-02, Vazyme Biotech, Nanjing, China) according to the manufacturer’s instructions. Signals were visualized using a CFX Connect Real-Time PCR Detection System (Bio-Rad, Hercules, CA, USA). The relative gene expressions levels were normalized to those of β-actin. Quantification was performed via the comparative 2^−ΔΔCt^ method. Primer sequences are displayed in Supplementary file 6.

### Western blot analysis

Total protein was extracted from cells or tissue by lysis with RIPA buffer (Cat# 89900, Thermo Fisher Scientific, USA) containing a protease inhibitor cocktail (Cat# 20-116, Millipore, St. Louis, USA) and phenylmethanesulfonyl fluoride (PMSF, Cat#ST506, Beyotime). Lysates were collected and centrifuged at 12000 rpm for 30 min at 4 ℃, and the protein concentration in the supernatant was determined using a BCA Protein Assay Kit (Cat# 23225, Thermo Fisher Scientific). Afterward, all the protein samples were normalized for protein concentration and separated by electrophoresis at a concentration of 20-30 μg on a 10% SDS-PAGE gel. Proteins in the SDS gels were transferred to PVDF (Cat# IPVH00010, Millipore) membrane using an electroblotting apparatus (Bio-Rad, Hercules, CA, USA). After 1 h of sealing with 5% skimmed milk, primary antibodies (See Supplementary file 5) were added and incubated overnight at 4 °C. Membranes were then rinsed five times (7 min each) with TBST and incubated with the corresponding horseradish peroxidase-conjugated secondary antibody at room temperature for 1 h. Membranes were rinsed five times (7 min each) with TBST, and immunoreactions were detected by enhanced chemiluminescence (ECL) detection and analyzed with the ImageJ software. The protein expression levels were normalized to β-actin.

### Transepithelial Electrical Resistance (TER) measurement

Transepithelial Electrical Resistance (TER) detected by using a Millicell-electrical resistance system (ERS)-2 Volt-ohm meter (Cat# MERS00002, Millipore) was used to evaluate the integrity of the Sertoli cell TJ-permeability barrier. Briefly, SCs were plated on transwell chambers (Corning; diameter: 6.5 mm; pore size: 0.4 μm; effective surface area: 0.3 cm^2^) at 1.0 × 10^5^ cells per well. Each transwell chambers was placed inside the well of a 24-well dish with 0.5 mL DMEM each in the apical and the basal compartments. Riluzole (10 μM) and/or BSF (200 μM) was supplemented in cell culture medium on day 2, the reaction lasted for 24 h. TER readings were recorded daily till day 4, the medium was freshly supplied daily.

### Molecular docking

The initial structure of riluzole was obtained from the PubChem database. The structure of riluzole is energy minimized by Chem3D and converted to mol2 format. Import small molecular compounds into AutoDock Tools software and add atomic charges, assign atom types, make all flexible bonds rotatable by default, and finally save them as pdbqt files. The 3D structure of TRPC3 (PDB ID: 5ZBG), TRPC5 (PDB ID: 7D4P), TRPC6 (PDB ID: 5YX9), and TRPC7 (PDB ID: 6MIX) were obtained from the Protein Data Bank database (https://www.rcsb.org/). After using Pymol 2.1 software to delete irrelevant small molecules in the protein molecule, import the protein molecule into the AutoDock Tools software, delete water molecules, add hydrogen atoms and set the atom type, and finally save it as a pdbqt file. The processed riluzole compound was used as a small molecule ligand, and the TRPC3/5/6/7 protein targets was used as a receptor. AutoDock Vina was used to completing the molecular docking. The center position of the Grid Box determined according to the interaction between the small molecule and the target (TRPC3: x= 127.95, y= 114.91, z= 156.02; TRPC5: x= 159.804, y= 122.164, z= 179.562; TRPC6: x= 128.02, y= 115.18, z= 155.95; TRPC7: x= 127.95, y= 114.91, z= 156.02). The width and height of grid box are both set to 30×30×30 Å. Pymol 2.1 software was used to visualize the binding effect of compounds and proteins.

### Statistical analysis

All experiments were repeated at least three times. All of the data were analyzed using GraphPad Prism software (Version 8.0, San Diego, CA, USA) and they were expressed as mean ± SEM. Data were compared by a twotailed unpaired Student’s t-test or one-way analysis of variance for multiple comparisons. Differences were considered significant at *P*<0.05 in all of the cases.

### Statement of ethics

All animal experiments were approved by the Institutional Animal Care and Use Committee (IACUC) of Jinan University (IACUC Approval: 20200327-68) and were conducted in accordance with the National Institutes of Health Guide for the Care and Use of Laboratory Animals.

## Data availability

The data supporting the findings of this study are available from the corresponding author upon request.

## Acknowledgements

This work was supported by the National Natural Science Foundation of China (No. U22A20277, 32170865, and 82071634), the Natural Science Foundation of Guangdong Province (No. 2022A1515012178), the Guangzhou Key R&D Program (No. 202103030003), and Guangdong Key Areas R&D Program (No.2022B1111080007).

## Author contributions

**Rufei Huang:** Conceptualization, Software, Methodology, Investigation, Data curation, Writing – original draft preparation. **Huan Xia:** Methodology, Data curation, Validation. **Wanqing Lin:** Software, Formal analysis. **Zhaoyang Wang:** Validation. **Lu Li:** Visualization. **Jingxian Deng:** Visualization. **Tao Ye:** Investigation. **Ziyi Li:** Investigation. **Yan Yang:** Conceptualization, Writing – review & editing, Supervision, Resources, Funding acquisition. **Yadong Huang:** Conceptualization, Writing – review & editing, Supervision, Resources, Funding acquisition.

## Supplementary file

**Supplementary file 1.** Riluzole administration enhanced sperm quality of oligospermic mice, related to Figure 1. (**A**) Body weight at week 8 in mice with oligospermia after riluzole treatment (n = 6). (**B-I**) Straight-line velocity (VSL), Curvilinear velocity (VCL), Average path velocity (VAP), Amplitude of lateral head displacement (ALH), Mean angular displacement (MAD), Beat-cross frequency (BCF), Straightness coefficient (STR), and Linearity (LIN) of sperms in oligospermic mice after administration of riluzole (n=6). The data were analyzed for more sperm counts over 1000 with CASA. Values are expressed as mean ± SEM. One-way analysis of variance (ANOVA) was used to analyze statistical differences; ***P < 0.001, **P < 0.01, *P <0.05 compared to BSF.

**Supplementary file 2.** Validation of testicular tissue culture model in vitro. (**A**) Schematic diagram of in vitro culture of oligospermia mice testis tissue. Busulfan-induced oligospermic mice were established in 4-week-old male mice, followed by euthanization in a benchtop and transfer of testes to 1.5% agarose gel blocks in 12-well plates. Complete medium containing different concentrations of riluzole (0.1 μM, 1 μM, 10 μM, and 50 μM) was added to half the height of the agarose gel blocks. The medium was changed every two days during the culture. 7 days after administration, the testes were fixed in 4% paraformaldehyde for preparation of paraffin sections and HE staining. (**B**) Histopathological analysis of testicular tissue in vitro. Black arrow presented low numbers of germ cells in the seminiferous tubule lumen. Scale bar, 100 μm.

**Supplementary file 3.** Riluzole had no effect on sexual hormone levels of oligospermia mice. (**A**) Testosterone concentration in serum of oligospermia mice after riluzole treatment (n=6). (**B**) LH concentration in serum of oligospermia mice after riluzole treatment (n=5). Values are expressed as mean ± SEM. One-way analysis of variance (ANOVA) was used to analyze statistical differences; \*\*\**P* < 0.001, \*\**P* < 0.01, \**P* <0.05 compared to BSF.

**Supplementary file 4.** Riluzole administration in oligospermia mice, related to Figure 1 and 2.

**Supplementary file 5.** Primary antibodies used in the experiments.

**Supplementary file 6.** Primer sequences used in qRT-PCR analysis.

